# Mechanosensitive interactions between Jag1 and Myo1c control Jag1 trafficking in endothelial cells

**DOI:** 10.1101/2025.03.27.645426

**Authors:** Oscar M.J.A. Stassen, Noora Virtanen, Kai-Lan Lin, Freddy Suarez Rodriguez, Matthijs J.M. Heijmans, Feihu Zhao, Garry L. Corthals, Carlijn V.C. Bouten, Cecilia M. Sahlgren

## Abstract

Morphogenesis of the cardiovascular system is responsive to hemodynamic cues. In endothelial cells the organization of morphogenic signaling proteins can be regulated by membrane presentation and internalization of proteins. How these signaling proteins are regulated by hemodynamics is unclear. One of the signaling proteins that is regulated by hemodynamics is Jagged1, a ligand in the Notch pathway. Here we set out to identify factors that differentially interact with Jagged1 in response to shear stress exposure, by combining an orbital shaker as a shear stress platform with endothelial cells expressing Jagged1 coupled to an APEX2-tag for proximity labeling. Myo1c was identified and confirmed through coimmunoprecipitation as a Jag1 interacting factor under static conditions, with reduced interaction after exposure to shear in endothelial cells. We showed that Jagged1 polarized downstream of shear followed by nucleograde transport of Jagged1. Myo1c knockout inhibited shear-induced Jagged1 polarization and consequent nucleograde transport. Further, Myo1c knockdown reduced membrane levels of Jagged1 under static conditions, but not under shear conditions. Together, our data reveal a role for Myo1c in the hemodynamic control of Jagged1 localization in endothelial cells.

## Introduction

Surface membrane proteins play a central role in mediating cellular interactions and signaling. To control interactions, cells continuously refine the protein population at their surface through regulated protein trafficking (Foot et al., 2017). The surface membrane protein Jagged1 (Jag1) is one of the ligands of the Notch receptor family and has important functions in the vasculature. Loss of Jag1 is embryonically lethal, and various congenital or acquired cardiovascular diseases are associated with Jag1 mutation or dysfunction, including atherosclerosis, Alagille syndrome, or Tetralogy of Fallot (Mašek and Andersson, 2017; Souilhol et al., 2020; Souilhol et al., 2022; Stassen et al., 2020).

Notch is a highly regulated and dose-sensitive signaling pathway (Bray, 2016). There are several mechanisms to control the dose of Notch activation, including posttranslational modifications, control of membrane presentation, and combinatorial levels of ligands of varying affinity presented in cis or in trans (Antfolk et al., 2019), but the mechanisms with which the cell controls trafficking of Jagged are poorly understood (Antfolk et al., 2017; Seib and Klein, 2021). In endothelial cells, Jag1 levels and their distribution within the endothelial network have been shown to be posttranscriptionally controlled by the ZFP36 mRNA decay protein (Sunshine et al., 2024), and the activity of Jag1 in Notch signal activation is intricately linked to the mechanical state of the cardiovascular system (Stassen et al., 2020; Suarez Rodriguez et al., 2023). Jag1 has been shown to have distinct molecular mechanical properties, possibly related to catch bond principles (Luca et al., 2017). At the cellular level, Jag1 reorganizes under shear stress, which promotes its functionality as a signal sender in Notch signaling (Driessen et al., 2018; Souilhol et al., 2020).

In this study, we optimized a proximity labeling approach in combination with a proteomics-compatible shear stress platform and identified Myo1c as a shear-stress regulated factor recruited to Jag1 in endothelial cells. In the absence of shear Myo1c interacts with Jag1, whereas in the presence of shear this interaction is reduced. Under shear stress Jag1 polarized downstream of the direction of shear. Knocking down Myo1c inhibited this shear-induced polarization and pharmacological inhibition of Myo1c reorganized Jag1 in perinuclear structures under static conditions. In addition, upon knockdown of Myo1c the membrane levels of Jagged1 were reduced under static conditions, whereas under shear conditions this effect trended towards an increase in membrane levels, indicating a complex interplay between Myo1c, shear stress and Jag1 trafficking. Together, these data reveal Myo1c as a novel factor responsible for trafficking of Jag1 under hemodynamic loading.

## Materials and methods

### Plasmids

To introduce Jag1-APEX2 into endothelial cells a pLenti Jag1-APEX2 lentivirus plasmid was constructed (see supplemental methods for plasmids and cloning). As control plasmid for the proximity labeling we used an APEX2 domain coupled to a nuclear export signal for cytoplasmic labeling. pcDNA3 APEX2-NES (RRID:Addgene_49386) was a gift from Alice Ting through Addgene.

### Cell Culture

Human Umbilical Vein Endothelial Cells (Lonza, pooled, Cat#C2519A) were cultured in Endothelial Cell Basal Media 2 (Promocell) supplemented with Growth Medium 2 supplement mix (composition in Supp. Methods). HEK293T, MCF7, and HeLa cells used for producing and testing lentivirus and PCLP assays were cultured in DMEM high glucose (Gibco, Cat#41966) supplemented with 10% fetal bovine serum, 100 units/ml penicillin, and 100 ug/ml streptomycin. All cell cultures were maintained at 37°C and 5% CO_2_ in a humidified incubator and passaged twice per week.

### Lentiviral production and transduction

Lentivirus was produced by transfecting pLenti transfer plasmid, pCMVR8.74, and pMD2.G into HEK293T cells using PEI transfection. Medium was harvested in the 3 days following transfection, precleared by 500 g centrifugation and filtered through a 0.45 um syringe filter. Filtered medium was centrifuged for 120 min at 50,000 g at 16°C and pellets were resuspended in 1x PBS for storage at - 80°C and transduction. pCMVR8.74 and pMD2.G were a gift from Didier Trono (RRID:Addgene_22036 and RRID:Addgene_12259).

### Mechanoresponsive Jagged Interacting Factor identification

Labeling and isolation of interacting factors through APEX2 were performed as described before (Hung et al., 2016). HUVECs were transduced with Lenti Jag1-APEX2, cultured at 17,000 cells / cm^2^ on Collagen IV coated annular culture vessels (Dish-in-a-Dish) of 13.5 cm outer diameter and 5.6 cm inner diameter (Driessen et al., 2020). Confluent cells were cultured under static (n=2) or shear conditions (15 min (n=1) or 24 hour (n=2), 150 rpm, 3 mm medium height, 1 cm orbit diameter, resulting in a shear between 0.6 and 0.8 Pa, and subsequently cultured for 30 mins under static or shear conditions in the presence of 500 uM biotin-tyramide. Biotin labeling was initiated by adding 0.1% hydrogen peroxide for 60 seconds after which the reaction was quenched by washing 3x with PBS supplemented with quencher (5 mM trolox, 10 mM sodium ascorbate and 10 mM sodium azide). Samples were then lysed in RIPA (50mM Tris, 150mM NaCl, 0.1% w/v SDS, 0.5% w/v sodium deoxycholate, 1% triton) supplemented with Complete protease inhibitor (PI; Roche) and quencher (RIPAPIQ). To isolate biotinylated proteins, magnetic streptavidin beads (Pierce) were equilibrated in RIPAPIQ. The samples were incubated with the beads under constant agitation for at least 4 hours or overnight at 4°C. Beads were washed two times with RIPA, subsequently with 1M KCl, then 0.1M Na_2_CO_3_, then 2M Urea in 10 mM TrisHCl, and twice with RIPA again. Reduction and alkylation occurred by incubation with 10 mM DTT in 100 mM Ammonium Bicarbonate and 100 ul of 55 mM iodoacetamide for 1 hour each, and a final incubation in 10mM DTT, 8 mM iodoacetamide in 100 mM ammonium bicarbonate. Finally, bead-bound products were cleaved with 1 ug mass spectrometry grade trypsin. Cleaved product was freeze-dried and analyzed by mass spectrometry on a TripleTOF5600 (Sciex). Sample Spectra were analyzed using MetaMorpheus version 0.0.303 to identify peptide spectrum matches.

### GO analysis

Gene enrichment analysis was conducted using g:Profiler, a web-based server (biit.cs.ut.ee/gprofiler/gost). The functional profiling of the enrichment results was carried out in g:Gost (version e101_eg48_p14_baf17f0) using the g:SCS algorithm with the significance threshold of 0.05. The most relevant GO terms were filtered and visualized.

### Total membrane protein biotinylation

Cells after static and shear treatment were placed on ice and washed three times with ice cold PBS-CM (0.5 mM MgCl2 and 0.9 mM CaCl2), followed by membrane protein labeling with 0.5 mg/ml EZ-link sulfo-NHS-SS-biotin (Thermo Scientific, 21331) in PBS for 30 min on ice. Afterwards, the cells were washed twice with ice-cold 0.1 M glycine in PBS, then twice with PBS-CM. The cells were lysed in RIPA+PI (RIPA buffer with cOmplete Protease Inhibitor Cocktail (Roche, 4693116001)) for 10 min on ice and centrifuged at 15,000 rpm for 10 min at 4°C. The supernatant was incubated in streptavidin agarose beads (Thermo Scientific, 20353) and incubated overnight at 4°C with a rotary shaker. The next day, samples were centrifuged at 14,000 rpm for 30 s and washed with RIPA+PI three times. The pellet was resuspended in Laemmli sample buffer for western blot.

### Western Blotting

Cells were washed with 4°C PBS and incubated with RIPA lysis buffer at 4°C for 20 min after which the contents of the wells were harvested. Samples were centrifuged at ≥10,000 g, at 4°C and the supernatant was retained. 4x Laemmli buffer was added to the supernatant and heated for 5 min at 95°C. SDS-Page (sodium dodecyl sulfate polyacrylamide gel electrophoresis) with precast gels (12% or 4-12% gradient gels; Biorad) were used to separate proteins. Proteins were transferred to a nitrocellulose membrane (Amersham, Protran). Both the separation and the transfer of the proteins was carried out in a wet transfer blotting system (Biorad) at 100 Volt. Membranes were blocked with 5% nonfat dry milk in PBS with 0.1% Tween, for 1h at room temperature. Primary staining on the membranes was performed overnight at 4°C. The membranes were then washed 3x in PBS 0.1% Tween and a secondary staining with HRP-coupled antibodies was done for 1h at room temperature. The membranes were washed at least 3x after secondary staining. Detection was performed with a ECL chemiluminescence kit on an iBright CL1500 (Invitrogen).

### Coimmunoprecipitation

Cells were lysed with lysis buffer (50 mM Tris (pH 7.5), 150 mM NaCl, 1% Triton X-100, 0.1% SDS, supplemented with protease and phosphatase inhibitors). Samples were cleared using a Branson sonifier three times for 5 seconds and spun down at 14k g. Beads were washed before use, three times with 50 mM Tris-HCl (pH 7.5); 250 mM NaCl; 0.1% NP-40. Samples were pre-cleared and allowed to conjugate to antibody overnight. Proteins were CO-IPed with magnetic beads and run on SDS-PAGE. Loading was measured using Revert 700 Total Protein Stain (LICOR).

### Shear stress live imaging

HUVECs transduced with Lenti Jag1-eGFP were sheared in a parallel plate setup (ibidi). This was placed on a Leica DMI8 microscope, with a Leica EL6000 as light source, a HC PL APO 40x/0.95 air objective, and a filter set of 450-490/500/500-550 for imaging eGFP. Cells were cultured at a density of 50,000 cells/cm^2^ in μ-slides VI 0.4, coated with collagen IV (ibidi, cat.# 80602) and connected to an ibidi pressure pump system in the incubation box of the microscope. At the start of an imaging experiment shear stress was built up linearly in 4 steps over 4 hours to the final desired shear stress of 1.8 Pa. Images were acquired with a Hamamatsu-C13440-20CU-US-300708 camera, every 4 minutes, with a 2s exposure for the Jag1-eGFP channel. Live recordings were processed to remove bright outlier (hot) pixels, by “remove outliers” in ImageJ (radius 2.0, threshold 100).

### Immunocytochemistry

Cells after static and shear treatment were fixed with 4% PFA for 10 min at room temperature, followed by permeabilization at room temperature with 0.1% Triton-X in PBS for 5 min. The cells were blocked for 1 hour at room temperature with 3% BSA with 0.05% Triton-X. The cells were incubated overnight at 4 °C with primary antibodies for Myo1c (1:50 dilution)(Santa Cruz Biotechnology, sc-136544) and Jag1 (1:100 dilution) (Cell Signaling Technology, #2620). The cells were washed three times with PBS before incubating with Alexa 488 anti-rabbit secondary antibody (1:500), Alexa 555 anti-mouse secondary antibody (1:500), and DAPI (1:1000) at room temperature for 1 h in the dark. The cells were washed twice with PBS and kept in PBS with 0.1% sodium azide in the dark, at 4 °C. Fluorescence imaging was performed using a confocal laser-scanning microscope (LSM880; Carl Zeiss).

### siRNA Knockdown and inhibitors

Knockdown of Myo1c was performed by transfecting 25 nM siRNAs into target cells using an RNAiMAX transfection protocol. To target Myo1c, 4 pooled siRNAs were transfected (L-015121-00; Dharmacon Reagents), and as control 4 pooled Non-Targeting siRNAs were transfected (D-001810-10, Dharmacon Reagents). Cells were incubated for 4 hours with the transfection mix and provided with fresh medium. For inhibition of Myo1c, samples were treated with 5 μM of pentachloropseudilin, with DMSO as vehicle control (PCLP; Aobious).

## Results

### Shear induces Jag1 reorganization into dynamic substructures downstream of shear

To study the reorganization of Jagged under shear, we exposed endothelial cells to shear stress and stained them for Jag1. In addition to the emergence of reorganized clustering of Jag1 under shear as described before (Driessen et al., 2020), polarization of Jag1 was observed downstream of the shear direction (Figure 1B). Early after shear onset Jag1 rapidly accumulates at the planar pole downstream of shear, followed by nucleograde transport from the membrane (Fig. 1C-E & Supp. Movie 1-3 (Shear) and Supp. Movie 4-5 (static). The live imaging showed that after polarization of Jag1, the endothelial cells initiate migration in the direction of shear. Under static conditions we rarely observed reorganization of Jag1, only in cases when a cell protrusion retracted.

**Figure 1.**
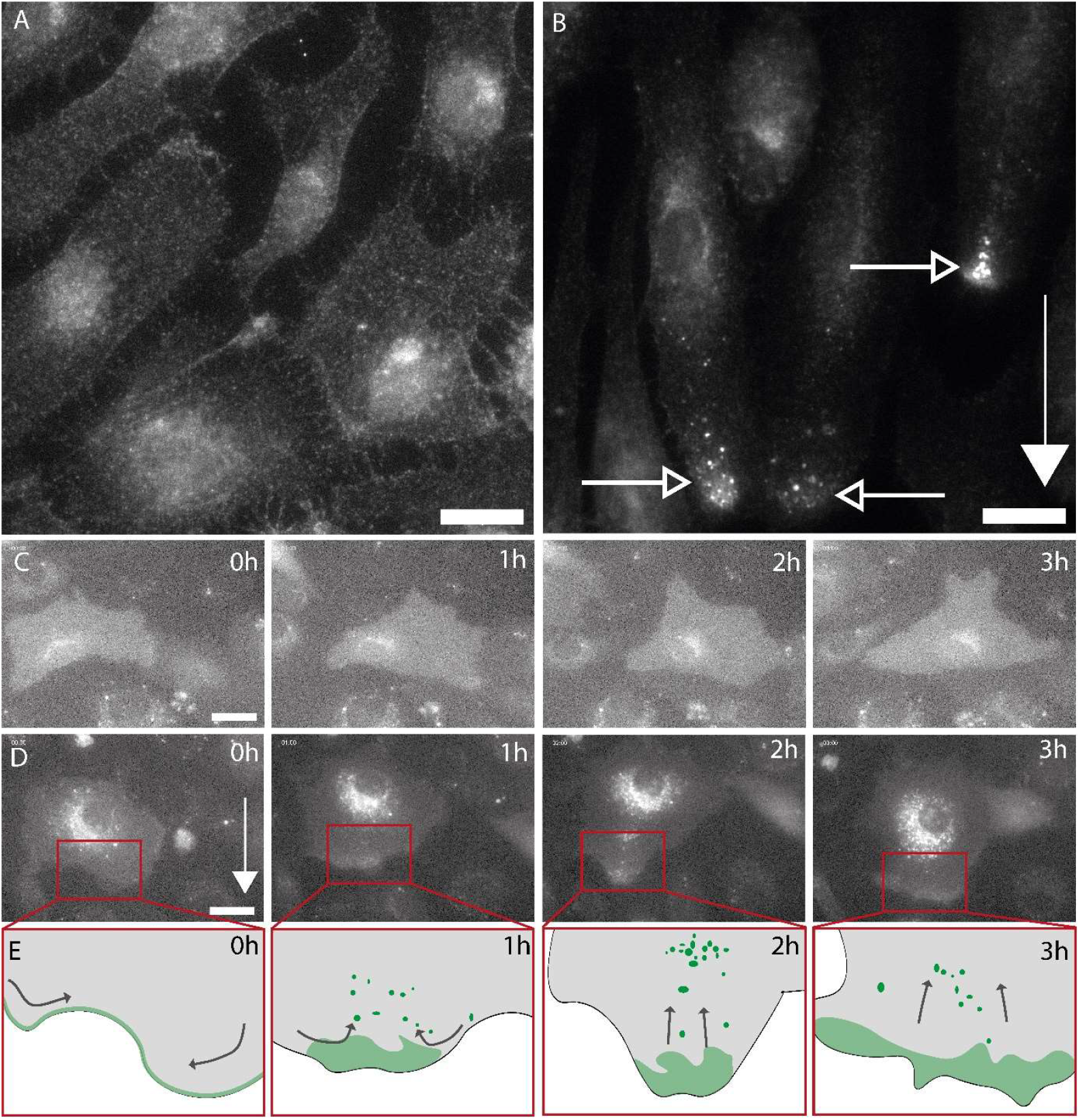
Jagged1 relocalizes in response to shear stress. A,B) Confocal microscopy images of Jag1 immunofluoresence staining. HUVECs exposed to static (A) and 2 Pa shear stress for 48 h (B) and samples were fixed and stained with a Jag1 antibody. The shear direction is indicated with filled arrows, and shear-induced Jag1 relocalization is indicated with open arrows. Scale bar 20 μm. See supplementary figure 1 for an expanded field of view of Fig 1B. C, D) Stills from live imaging of Jag1-eGFP expressing HUVECs under static (C) or shear (D). Scale bar 25 μm. See Supplemental movies 1-5 for recordings at 4-minute intervals. E) Manual traces of the cell outline (gray) and shear induced Jag1 localization (green) observed in the magnification windows (red). Black arrows indicate the direction the Jag1 densities appear to follow.

### Jag1-APEX2 identified shear-sensitive interactors

To identify mechanoresponsive Jag1-interacting factors that contribute to Jag1 trafficking, we took a combined approach of a large surface area shear system and proximity labeling (Driessen et al., 2020). We genetically attached the engineered ascorbate peroxidase (APEX2) to Jag1. APEX2 labels nearby peptidesin the presence of hydrogen peroxide and biotin-tyramide (Fig. 2A) (Hung et al., 2017). Tagging the C-terminal intracellular domain of Jag1 enabled labeling of proximal proteins (within a 20 nm radius) in the presence or absence of flow-induced shear stress (Fig. 2B). Verification of the construct showed that Lenti-Jag1-APEX2 exhibited a higher molecular weight than endogenous Jag1, as expected (Fig 2C). The localization of the Jag1-APEX2 or NES-APEX2 construct was tested by incubating transduced cells with 500 μM biotin-tyramide for 30 mins and subsequently adding 1 mM of hydrogen peroxide for 1 minute before quenching the reaction. This showed the expected localization of labeling patterns, NES-APEX2 having enriched cytoplasmic labeling whereas Jag1-APEX2 had enriched labeling at the membrane (Fig. 2D-E). As the shear stress platform, we utilized our previously modeled annular orbital shaker (or Dish-in-a-Dish) system with more defined and uniform shear characteristics than a simple dish on a shaker (Fig. 2G) (Driessen et al., 2020). To compare static and shear conditions, HUVECs expressing Jag1-APEX2 were cultured statically or exposed to a shear stress between 0.6 and 0.8 Pa for 15 min or 24h. Brightfield microscopic images of the cells during the labeling experiment confirmed shear-induced cellular alignment (Fig. 2H) (Driessen et al., 2020). The hits from the different shear exposure regimes that were labeled and are candidates for Jagged1 proximity are represented in Figure 3. The genes under each condition were subjected to GO analysis (Supp. Fig. 2). This analysis revealed proximity of Jag1 to proteins involved in membrane processes, adhesion, and the actomyosin network. Since the Myo1 family of motor proteins is involved in establishing cortical and membrane tension (McIntosh and Ostap, 2016), and Myo1c specifically has roles in regulating the presentation of membrane proteins and the stability of cell-cell adhesion (Tiwari et al., 2013; Tokuo and Coluccio, 2013), and recently was shown to augment shear-induced secretion of VWF from Weibel-Palade bodies (El-Mansi et al., 2024), we further investigated the Jag1-Myo1c interaction.

**Figure 2.**
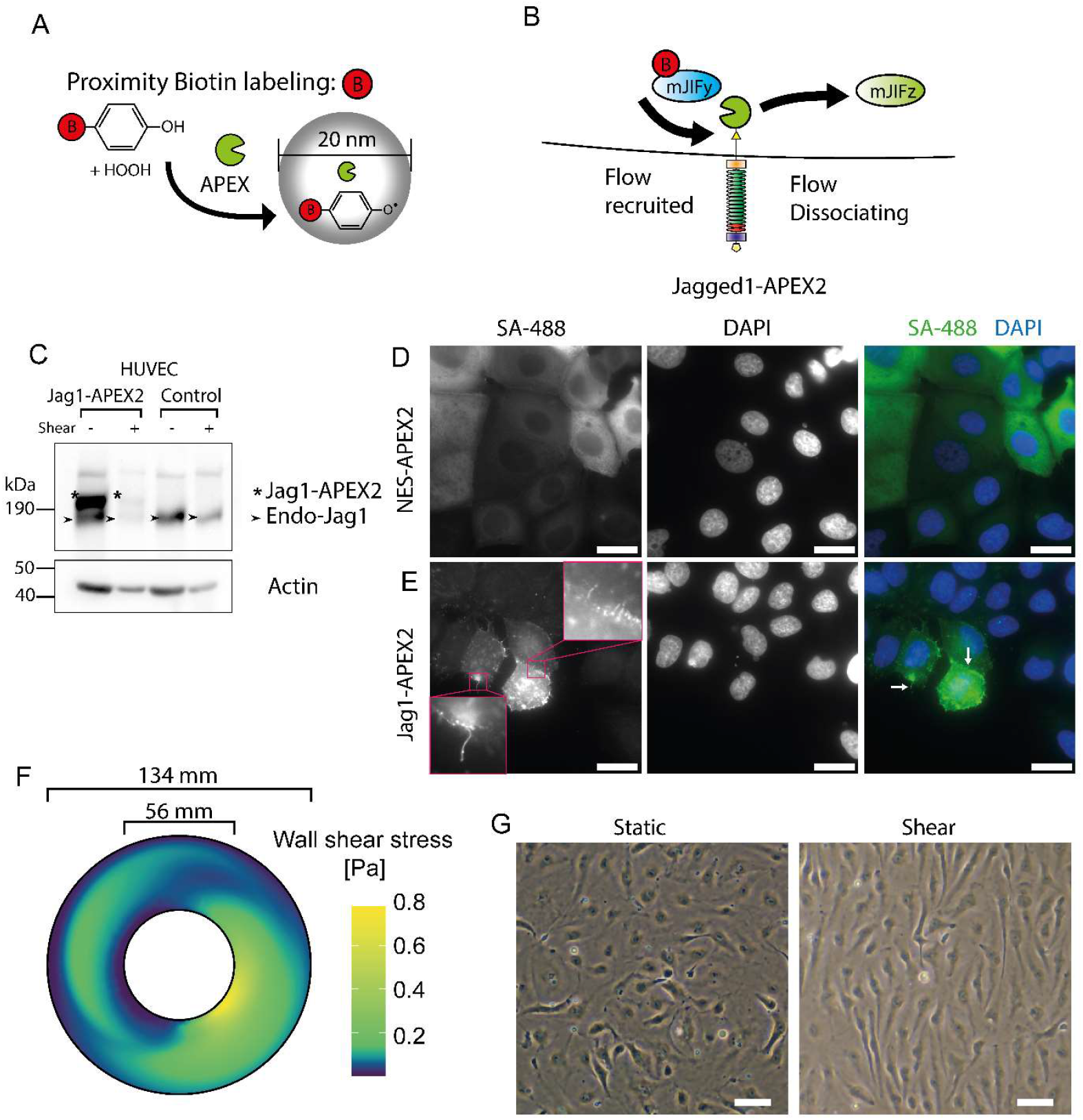
Proximity biotin labeling, mass spectroscopy and shear platform to identify shear stress sensitive Jag1 interacting proteins. A) The engineered ascorbate peroxidase (APEX2) convert biotin-tyramide to a free radical carrying species in the presence H_2_O_2_, allowing labeling of nearby peptides. Coupling APEX2 to Jag1 and performing labeling interaction under different mechanical loading conditions allows the identification of mechanoresponsive Jagged interacting factors (mJIFy & mJIFz). B) Jag1 engineered with APEX2 on the C-terminal (intracellular) domain enables labeling of cytoplasmic proximal proteins in the presence or absence of flow induced shear stress. C) Immunoblot of Jag1 and Lenti Jag1-APEX2 expression. HUVECs were transduced with Lenti-Jag1-APEX2, exposed to static or shear conditions, and blotted for Jag1 and actin. D-E) Immunofluorescence microscopy of cells labeled with streptavidin 488 (SA488, green) and DAPI (blue). MCF7 cells transduced with D) Lenti-APEX2-NES or E) Lenti-Jag1-APEX2, biotinylated and stained with streptavidin 488 and DAPI, scale bars = 25 μm. F) Visualization of computational model of the shear stress calculated for the annular culture platform (134 mm outer diameter, 56 mm inner diameter). G) Brightfield microscopy images of cells cultured statically (left) or under orbital shaker induced shear (right), scale bars = 50 μm.

**Figure 3.**
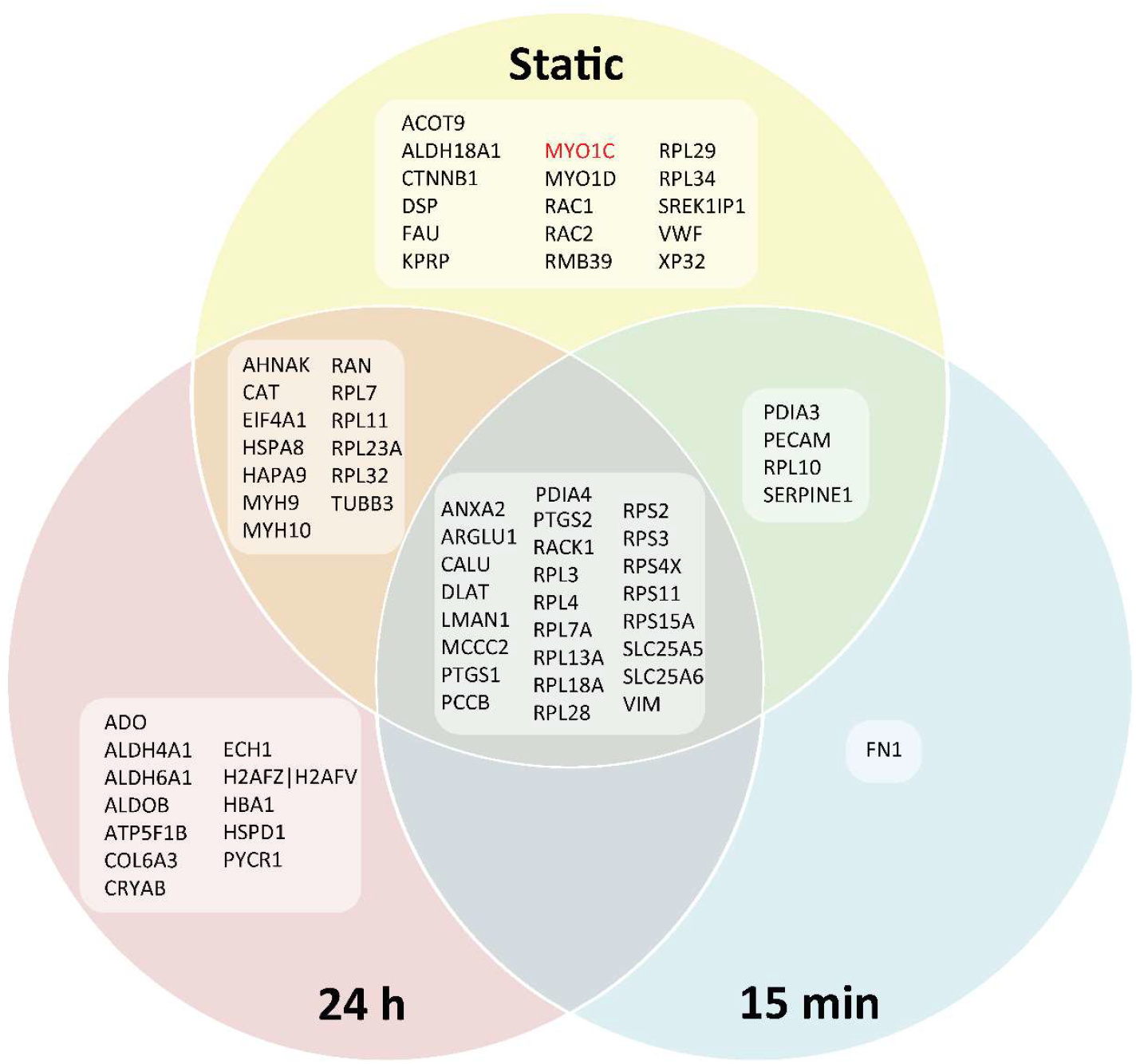
Biotin proximity labeling combined with mass spectrometry analyses reveal mechanosensitive Jag1 interacting proteins. Venn diagram with names of identified proteins. Represented are Jag1 interacting samples that returned a detection event in mass spec analysis in three different conditions: static, 15 min shear and 24 hour shear.

### Myo1c interacts with and regulates Jag1 localization

Myo1c emerged as proximal to Jag1 under static conditions, but was no longer detected when exposed to shear. Myo1c could effectively be knocked down in HUVECs and protein levels were not influenced by shear application (Fig. 4A), nor by knockdown of other hits identified in the screen. Under static conditions Myo1c coimmunoprecipitated (coIP) with Jag1, whereas under shear (0.8 Pa for 24h) this coIP was reduced (Fig. 4B&C). It was also possible to coIP Jag1 with a Myo1c pulldown (Supplementary Figure 4). To inhibit Myo1c we used PCLP, a noncompetitive reversible inhibitor of Myosin 1 class proteins, that prevents nucleotide binding by sterically hindering the coupling to the actin-binding site (Chinthalapudi et al., 2011). Treating HUVECs with an inhibitor of Myo1c, PCLP, caused a perinuclear accumulation of Jag1 within 2 hours of exposure, that further accumulates during exposure (Fig. 4D). This effect also occurred in cervical cancer HeLa cells (Supp. Fig. 3). As effects of PCLP have been reported on the cytoskeleton which might affect cellular organization, both HeLa cells and HUVECs were stained for actin, microtubules and intermediate filaments (vimentin), but no changes matching the Jag1 reorganization were visible, although a meshwork like organization of microtubules and vimentin appeared under PCLP treatment (Supp. Fig. 3C). After exposure to shear, both Jag1 and Myo1c polarized downstream of flow (Fig. 4E). Knocking down Myo1c using siRNA and exposing HUVECs to shear revealed that the flow-induced polarization was lost (Fig. 4F). Biotinylation of membrane proteins followed by streptavidin pulldown revealed that Jag1 membrane levels were reduced under shear (Fig. 4G & 4H). Knocking down Myo1c significantly reduced Jag1 membrane levels under static conditions. This reduction was not observed under shear; on the contrary, in the Myo1c knockdown shear induced a trend for increase of Jag1 membrane levels.

**Figure 4.**
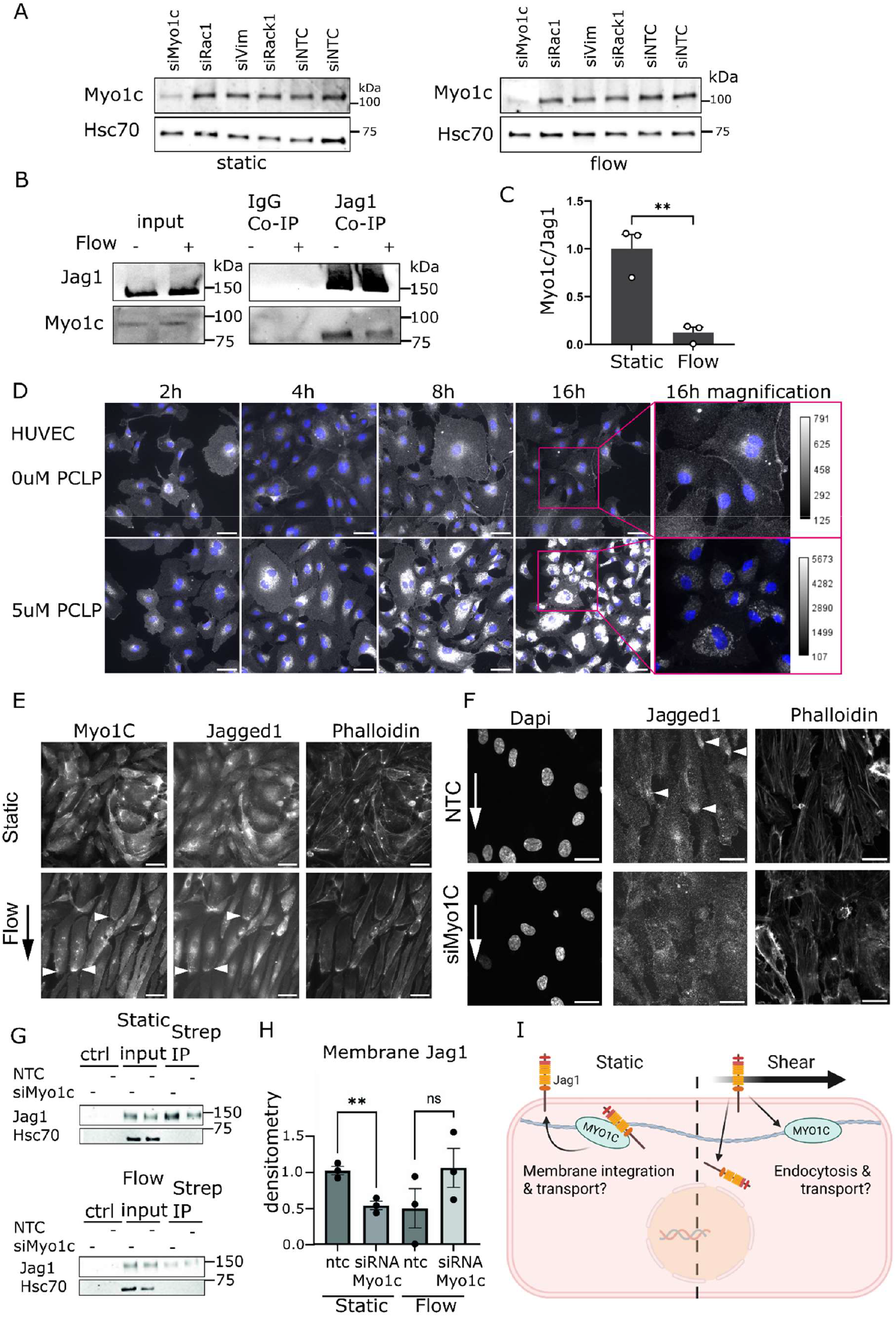
Myo1c interacts with and regulates Jag1 localization under shear stress. A) Immunoblot analysis of Myo1c expression. HUVECs were exposed to static or shear conditions in a swirling well, and lysates were analyzed for Myo1c levels with different Jag1 interacting candidate knockdowns. B) Jag1 and Myo1c co-immunoprecipitation. HUVECs cultured under shear (0.8 Pa for 24h) and static condition were lysed and immunoprecipitated using a anti-Jag1 antibody. The precipitate was immunoblotted for Myo1c. C) Quantification of B. D) Fluorescence microscopy images of HUVECs treated with PCLP and stained for DAPI (blue) and Jag1 (white). Figures in D from 2H up to 16H are represented with same lookup table (LUT), magnified images on the right were scaled with LUT tables as indicated to illustrate perinuclear Jag1 organization under PCLP treatment. Scale bars 50μm. E) Fluorescence microscopy images of HUVECs exposed to shear in parallel plate channels, stained for Myo1c, Jag1 and Actin (phalloidin). Flow-induced polarization of Myo1c and Jagged1 indicated by white arrowheads, scale bars 50 μm. F) Confocal fluorescence microscopy of HUVECs under parallel plate shear or static culture. HUVECs were treated with Myo1c siRNA or Non Targeting Control (NTC) and stained for Actin and Jag1. Scale bars 20 μm. G) Immunoblot of membrane proteins of HUVECs treated with siMyo1c or NTC, under static conditions and shear (200 RPM, 0.8 Pa) stained for Jag1 and HSC70. ctrl = Nonbiotinylated control, Strep IP = streptavidin pulldown. H) Quantification of F. I) Schematic representation of possible Myo1c and Jag1 interaction in the presence and absence of shear.

## Discussion

Our study reveals a novel role for Myo1c as a Jag1 shear-responsive motor protein. Myo1c, a member of the Myosin1 family, plays crucial roles in membrane tension and various cellular functions, including nuclear myosin activity and adaptation in vestibular cells (Batters et al., 2004; Gillespie, 2004; Venit et al., 2020).

In endothelial cells, Myo1c is vital for the membrane presentation of VEGFR2, G-actin transport, integrin trafficking (Cota Teixeira et al., 2019; Tiwari et al., 2013) and secretion of VWF (El-Mansi et al., 2024). Myo1c regulates protein delivery to and uptake from the plasma membrane under specific signaling cues, as is the case for Glut4 glucose transporter under insulin signaling in brown fat (Boguslavsky et al., 2012), and E-cadherin uptake and dynamics in epithelial cells (Tokuo and Coluccio, 2013). Our data suggest that Myo1c regulates bidirectional Jag1 trafficking and likely plays a role in Jag1 delivery to the plasma membrane under both static and shear conditions, and in Jag1 endocytosis and relocalization under shear (Fig 4E, G, H, I) (Driessen et al., 2018).

Only a few cytoplasmic tail interactors of Jag1 are currently known. A recent study on the nuclear Jag1 intracellular domain identified several candidate interactors in our screen, including PDIA4 and ANXA2 (Kim et al., 2022). Other interactors include Mindbomb and Neuralized, both RING ubiquitinases that promote the binding of Epsin and facilitate Jag1 endocytosis and Notch transactivation (Langridge and Struhl, 2017; Lee et al., 2015). We previously identified Vimentin as a Jag1 interactor and demonstrated that depletion of Vimentin increased membrane levels of Jag1 but decreased turnover and signaling (Antfolk et al., 2017; Engeland et al., 2019).

The mechanisms by which Myo1c alters Jag1 organization and its effects on signaling under shear stress require further exploration. Myo1c facilitates signal-dependent membrane protein presentation, such as VEGF-A-induced VEGFR2 presentation (Tiwari et al., 2013). Its absence reduces VEGFR2 on the plasma membrane, underscoring its role in angiogenesis regulation through modulation of VEGFR2 (Tiwari et al., 2013). Jag1 also regulates angiogenesis, (Antfolk et al., 2017; Benedito et al., 2009) suggesting additional roles of Myo1c in angiogenesis. Our screen also identified PECAM1 as a potential Jag1 interactor, forming a mechanotransduction complex with VEGFR2 (Coon et al., 2015). Further investigations on the role of Jagged in this context, especially its polarization under shear stress, is essential. Recent findings indicate that polarized mechanosensitive signaling domains can protect arterial cells from inflammation (Hong et al., 2023), suggesting Jag1 reorganization and its sensitivity to oscillatory shear in aortic cells may play a role in inflammatory signaling in the endothelium (Souilhol et al., 2020).

## Limitations of the study

Our study has certain limitations. The APEX2 tag used to identify targets binding to Jag1 was attached to the Jag1 protein through a linker to the C-terminal domain, where a PDZ ligand is located. Possibly, this prevented factors that bind through the PDZ domain from interacting with Jag1. Another possible mechanism of mechanosensitivity was recently also highlighted by the discovery of mechanosensitivity of the Afadin PDZ domain (Vachharajani et al., 2023). Afadin is one of the interactors of Jag1 through its PDZ-binding domain (Hock et al., 1998). In our screen, the weak and transient Afadin interaction, if relevant in endothelial cells, could not be detected.

Notably, while endothelial cells are generally described to migrate against the direction of shear, our observations of migration along the shear direction could arise from experimental conditions, such as growth factor concentrations and shear magnitude (Park et al., 2021; Vion et al., 2021). Determining how sensitive the regulation is to dynamic changes in shear will be an important future pursuit.

## Author Contributions

Conceptualization: OMJAS and CMS

Methodology: OMJAS

Software: FZ

Investigation: OMJAS, NV, KL, FSR, MJMH, GC

Formal analysis: OMJAS, KL, NV, FSR

Resources: CVCB and CMS

Data curation: OMJAS

Writing – original draft: OMJAS and CMS

Writing – review & editing: OMJAS, NV, KL, FSR, MJMH, FZ, GC, CVCB, CMS

Visualization: OMJAS

Supervision: CVCB and CMS

Project administration: OMJAS and CMS

Funding acquisition: CVCB and CMS

## Acknowledgements

This project has received funding from the European Research Council (ERC) under grant agreement number 771168 (ForceMorph) and the Academy of Finland under decision numbers 307133, 316882 (SPACE) and 330411 (SignalSheets). The research has also been supported by the InFLAMES Flagship Programme of the Academy of Finland (decision number 337531) and the Åbo Akademi University Foundation’s Center of Excellence in Cellular Mechanostasis (CellMech). We’d like to extend our gratitude to Tim Wezeman for preparing plasmids during Covid19-restricted lab-access, as well as to Iida Laiho, Jaakko Ahlberg, Meike Hulleman, and Anouk van der Net for help screening initial candidate interactors. We thank Peter Paul Franssen for producing a batch of Biotinyl-tyramide. We thank the Turku Bioscience center for discussion of Proteomics analysis and for imaging at the Cell imaging and Cytometry Core. pLenti PGK Puro DEST (w529-2) and pENTR1A-GFP-N2 (FR1) were gifts from Eric Campeau & Paul Kaufman (RRID:Addgene_19068 & RRID:Addgene_19364) – cDNA3 APEX2-NES was a gift from Alice Ting (RRID:Addgene_49386)).

