## Supplemental Information for "Mechanosensitive interactions between Jag1 and Myo1c control Jag1 trafficking in endothelial cells"

**Supplemental methods:**

**Plasmid generation**

pcDNA3 Jag1-APEX2-mKO2 was synthesized as the ORF from NM_000214.2 (Baseclear, Leiden, NL), coupled with a GDPPVAT linker to the APEX2 ORF including FLAG-tag from the Alice Ting lab (Lam et al., 2015), a TRTRPLE linker, and the mKO2 sequence (Sakaue-Sawano et al., 2008). To target HUVEC, a lentiviral variant without mKO2, due to size restrictions, was made by PCRing JAG1-APEX2 with the primers FW:TATTATCTAGATTATTTGTCATCATCGTCCTTATAATCGGCG and RV:CGAATGGAGTACATCGTAGGGGATCC, and cloned with XbaI and BamhI into pENTR1a (addgene #19364), and subsequently transferred to pLenti-DEST (Addgene #19068) using gateway recombination (LR-clonase 2).

The transfer plasmid for live imaging Jag1, pLenti Jag1-eGFP, was created through replacing the APEX2 in pENTR-Jag1-APEX2 with eGFP and recombination through the gateway method. Plasmids were expanded in NEB Stable and isolated using standard miniprep protocols (Qiagen), or midiprep protocols (Nucleobond).

**Medium growth factor composition**

Endothelial cells were cultured in Endothelial Cell Media with Growth medium 2 supplement mix. This consists of the following supplements and final concentrations: Fetal Calf Serum (0.02 ml/ml), Epidermal Growth Factor (recombinant human) (5 ng/ml), Basic Fibroblast Growth Factor (recombinant human) (10 ng/ml), Insulin-like Growth Factor (Long R3 IGF, recombinant human) (20 ng/ml), Vascular Endothelial Growth Factor 165 (recombinant human) (0.5 ng/ml), Ascorbic Acid (1 µg/ml), Heparin (22.5 µg/ml), Hydrocortisone (0.2 µg/ml).

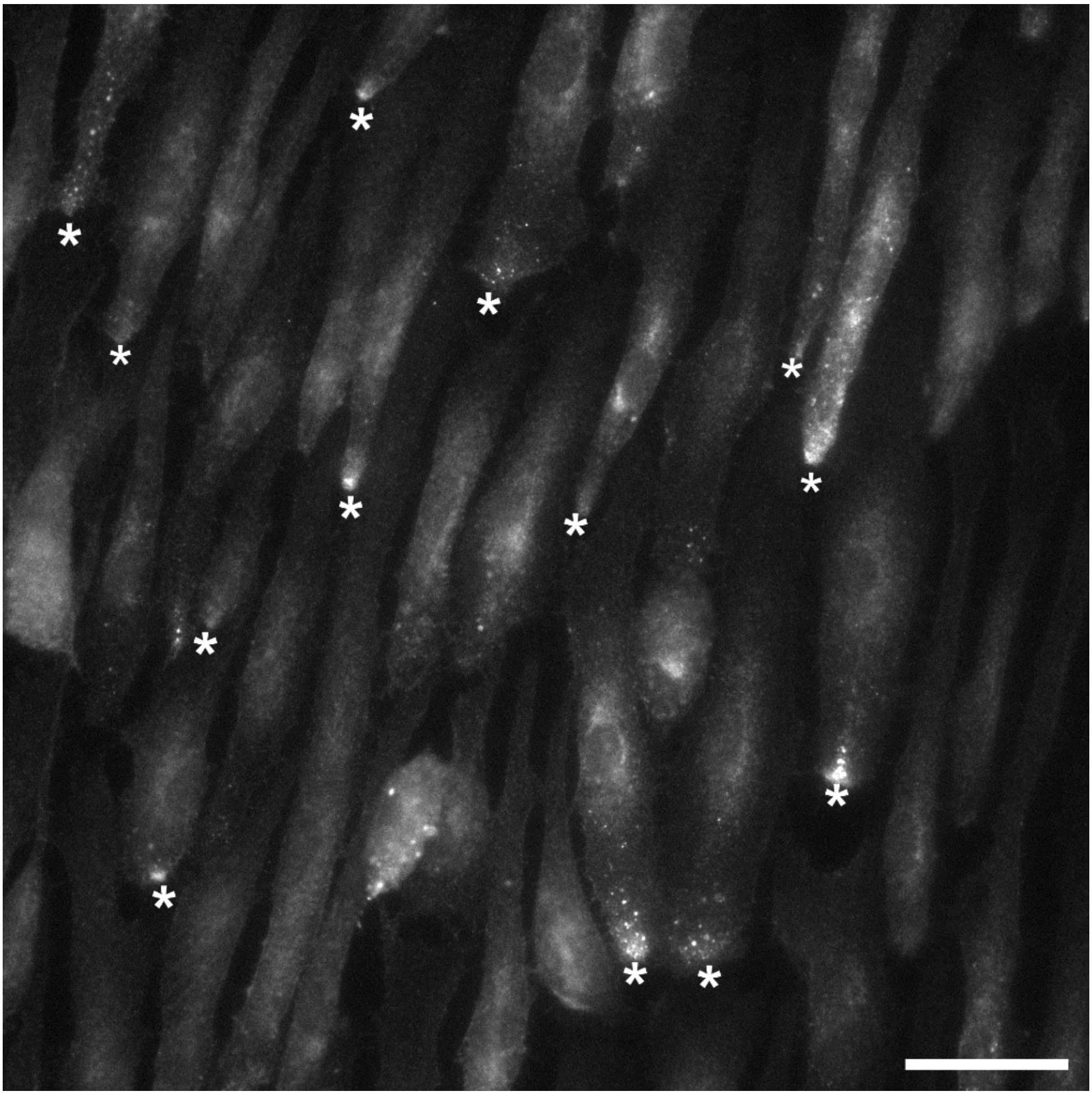

**Supplemental figure 1**

Fluorescence microscopy of larger field of view of cells in figure 1B. HUVECs were exposed to 48h of 2 Pa shear in a parallel plate chamber, fixed and immunostained for Jag1. Cells accumulate Jag1 downstream of shear, with the nucleograde chain of Jag1 between the pole and the nucleus visible.

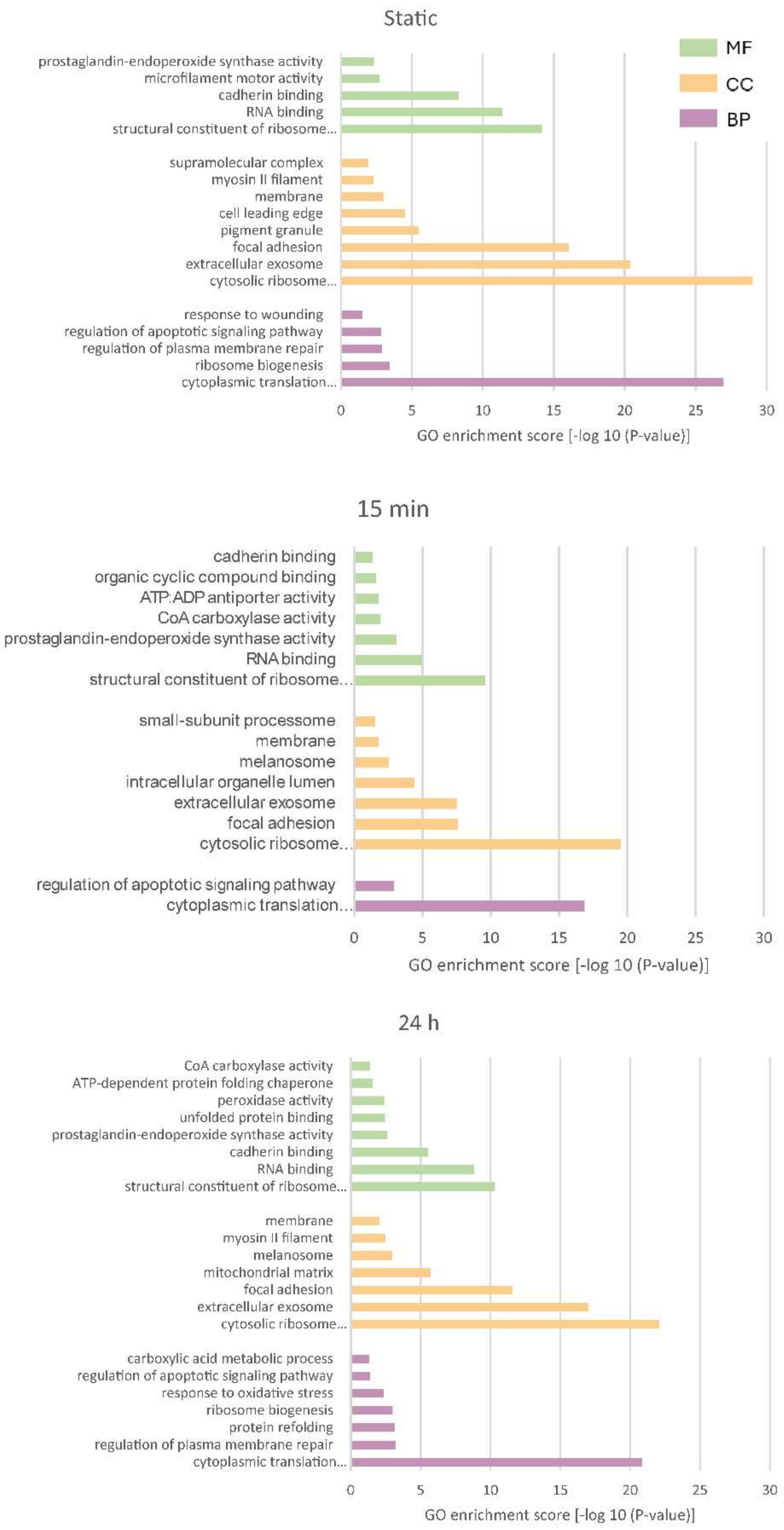


**Supplemental Figure 2**

Gene ontology analysis of Jag1-APEX2 identified proximal proteins. Analysis was performed for the static, 15 min shear and 24 hour shear samples. The Gene Ontology clusters Biological Process (BP; green), Cellular Compartment (CC; yellow) and molecular Function (MF; purple) were included.

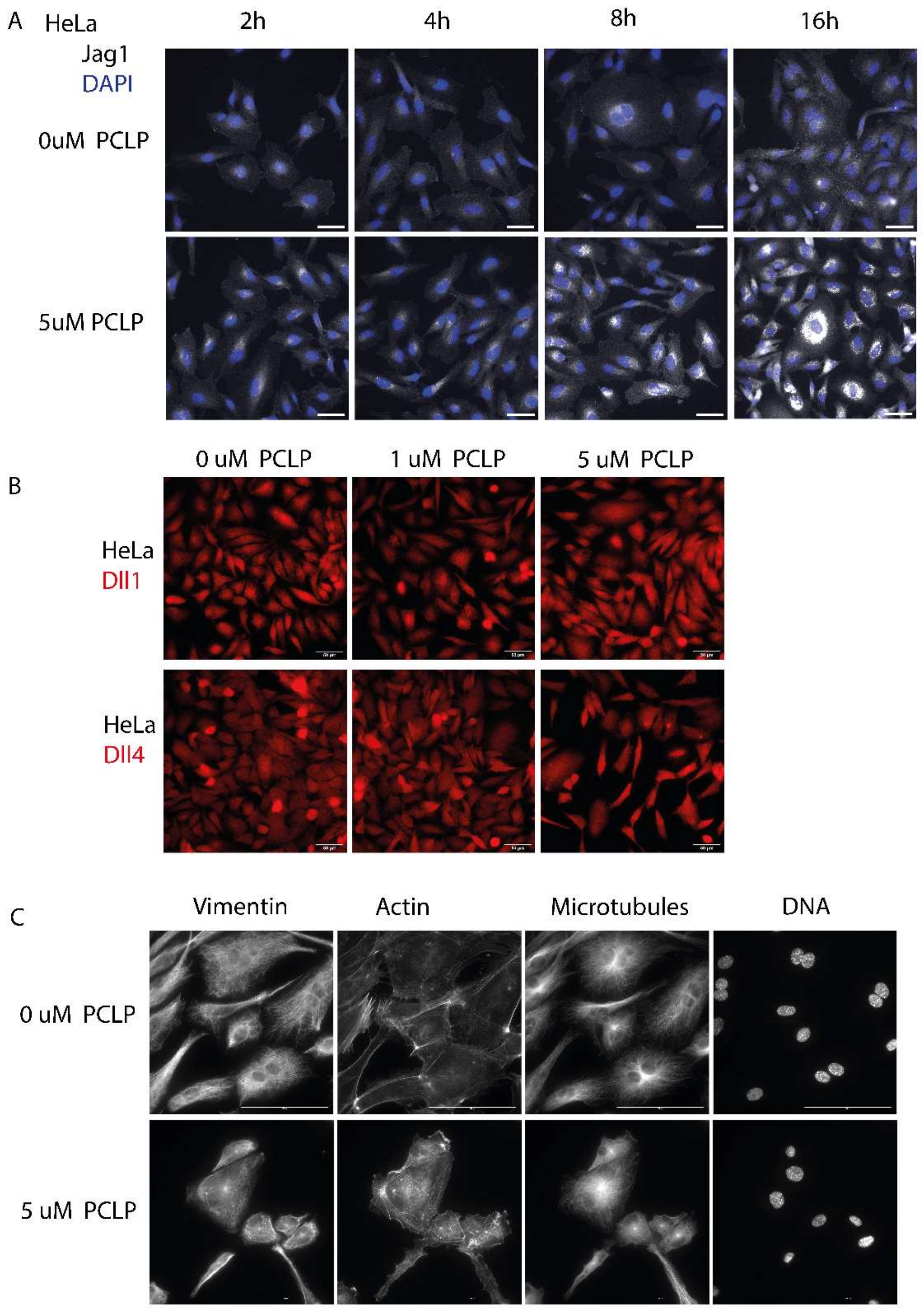


**Supplemental Figure 3**

Immunofluorescence microscopy images of A) HeLa cells were exposed to PCLP for various times. Reorganization of Jag1 did occur, but slower than in HUVEC cells. B) Specificity of the Jag1 response to PCLP was tested by staining DLL1 and DLL4 after exposure to PCLP. DLL1 and DLL4 did not show reorganization. C) HUVEC cytoskeleton after overnight exposure to 5 uM PCLP did not show the same alteration in organization as Jag1, changes in the actin cytoskeleton were observed and have been described before (Cota Teixeira et al., 2019).

**Supplemental Figure 4**

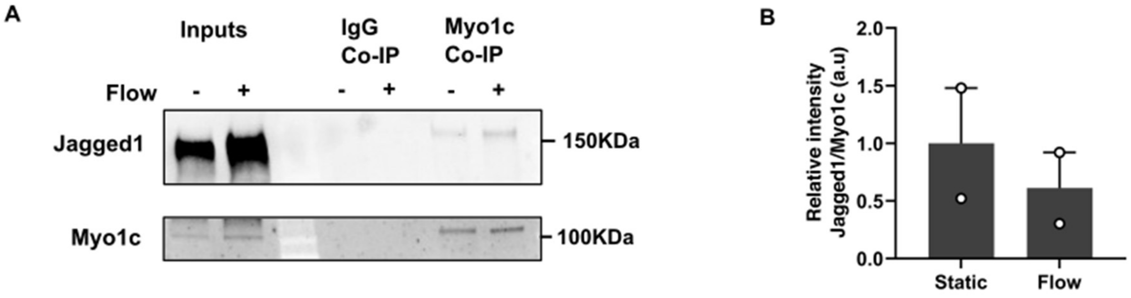


A) Jag1 and Myo1C co-immunoprecipitation. HUVECs cultured under shear (0.8 Pa for 24h) and static condition were lysed and immunoprecipitated using an anti-Myo1c antibody. The precipitate was immunoblotted for Jag1. B) Quantification of A.

**Supplemental Movies**

Live recordings of HUVECS stably transduced with Jag1-eGFP under shear (2Pa) (shear direction from top to bottom, Movies 1-3) or under static conditions (Movies 4 & 5). Scale bar = 20 µm.
